# A joint embedding of protein sequence and structure enables robust variant effect predictions

**DOI:** 10.1101/2023.12.14.571755

**Authors:** Lasse M. Blaabjerg, Nicolas Jonsson, Wouter Boomsma, Amelie Stein, Kresten Lindorff-Larsen

## Abstract

The ability to predict how amino acid changes may affect protein function has a wide range of applications including in disease variant classification and protein engineering. Many existing methods focus on learning from patterns found in either protein sequences or protein structures. Here, we present a method for integrating information from protein sequences and structures in a single model that we term SSEmb (Sequence Structure Embedding). SSEmb combines a graph representation for the protein structure with a transformer model for processing multiple sequence alignments, and we show that by integrating both types of information we obtain a variant effect prediction model that is more robust to cases where sequence information is scarce. Furthermore, we find that SSEmb learns embeddings of the sequence and structural properties that are useful for other downstream tasks. We exemplify this by training a downstream model to predict protein-protein binding sites at high accuracy using only the SSEmb embeddings as input. We envisage that SSEmb may be useful both for zero-shot predictions of variant effects and as a representation for predicting protein properties that depend on protein sequence and structure.

## Introduction

Small changes in the amino acid sequence of a protein can have a wide range of effects on its molecular structure, stability and function. Discerning the magnitude and consequences of such effects is central to understanding the molecular mechanisms of evolution and human disease (***Fowler et al., 2023***). Furthermore, the ability to manipulate sequences to change or optimize function fundamental to the field of protein engineering and design (***Freschlin et al., 2022***).

Decreased cost of DNA sequencing has enabled the development of new experiments that can generate biological data at an unprecedented scale. An example of this is the development of high-throughput assays that can provide a quantitative read-out of changes in activity, stability or abundance for thousands of protein variants in a single experiment. Such high-throughput assays, often called Multiplexed Assays of Variant Effects (MAVE) or Deep Mutational Scanning experiments, have enabled a substantial increase in the available data mapping the relationship between protein sequence and function (***Kinney and McCandlish, 2019; Rubin et al., 2021; Tabet et al., 2022; Notin et al., 2022a; Fowler et al., 2023***).

Many different types of MAVEs have been developed to probe different aspects of the protein sequence-function relationship. In particular, changes in protein abundance have been shown to be an important driver of change in protein activity (***Cagiada et al., 2021; Høie et al., 2022***). Thus, a specific type of MAVE called Variant Abundance by Massively Parallel Sequencing (VAMP-seq) has been developed to quantify variant effects on cellular protein abundance (***Matreyek et al., 2018***). By combining data generated by multiple MAVEs that probe different effects for each substitution—for example on abundance, activity or binding—it is possible to obtain mechanistic insights into how and why particular amino acid substitutions affect protein function (***Cagiada et al., 2021; Chiasson et al., 2020; Faure et al., 2022; Cagiada et al., 2023***).

Although MAVEs in principle can provide a complete mapping of the protein sequence-function relationship, the assays themselves are often costly and time-consuming. Furthermore, it is diffcult to design assays that probe all relevant functions of a protein and it may not always be possible to probe all possible variants in a single assay. The combinatorial explosion of multiple variants presents an additional challenge in experimental screening. In contrast, computational predictors of variant effects are able to make predictions for variants that have not been assayed before at a close-to-zero marginal cost and can serve as an additional source of information in the protein sequence-function mapping (***Fowler et al., 2023***).

Machine learning-based methods have proven to be a useful tool for predicting complex relationships in biology. Often, machine learning models are trained in a supervised manner, where the algorithm is trained to learn the mapping between a set of related input and target values. Although generally effective, such supervised learning algorithms can struggle to accurately predict the variant effects probed by MAVEs, possibly due to experimental differences between assays that make it diffcult to compare and standardize read-outs. In contrast, self-supervised algorithms that are trained directly on large amounts of input data only have emerged as a compelling alternative in the field of variant effect prediction (***Balakrishnan et al., 2011; Riesselman et al., 2018; Frazer et al., 2021; Meier et al., 2021; Pucci et al., 2022; Notin et al., 2022b; Cheng et al., 2023; Diaz et al., 2023***).

The majority of self-supervised predictors of variant effects rely on a single type of protein data that is used to learn the implicit correlations within the data. The output of a self-supervised predictor is often a probability distribution over the possible amino acid substitutions at a particular position in the protein. Examples of the types of data used as input include the wild-type amino acid sequence (***Lin et al., 2022; Brandes et al., 2022***), a multiple sequence alignment (MSA) (***Ng and Henikoff, 2001; Balakrishnan et al., 2011; Lui and Tiana, 2013; Nielsen et al., 2017; Hopf et al., 2017; Riesselman et al., 2018; Laine et al., 2019***) or the protein structure (***Boomsma and Frellsen, 2017; Jing et al., 2021a; Hsu et al., 2022***). Some methods have combined predictions from multiple protein data types at an aggregate level (***Strokach et al., 2021; Høie et al., 2022; Cagiada et al., 2023; Nguyen and Hy, 2023***), although some results suggest that a richer representation might be learned by combining multiple data types at the input level (***Mansoor et al., 2021; Wu et al., 2023; Wang et al., 2022; Yang et al., 2022; Chen et al., 2023; Cheng et al., 2023; Zhang et al., 2023***).

Here, we present the SSEmb (Sequence Structure Embedding) model (Fig. 1). The idea behind our model is that mapping multiple sources of protein information — in this case an MSA and the three-dimensional structure — into a single learned embedding should yield a model that is able to make more robust predictions. Specifically, we base our model on the MSA Transformer model (***Rao et al., 2021***), which can be used to make predictions of variant effects using a subsampled MSA as input (***Meier et al., 2021***). Although the MSA Transformer performs well in variant effect prediction, the accuracy of the predictions have been shown to be sensitive to the depth of the MSA used as input (***Notin et al., 2022a***). By constraining the MSA Transformer with structure information and combining the learned embeddings with a structure-based graph neural network (GNN) (***Jing et al., 2021b***), we show that we can obtain improved variant effect predictions when the input MSA is shallow. In contrast to other recent methods (***Chen et al., 2023***), SSEmb is trained fully end-to-end, with a focus on integrating and aligning sequence- and structure-based information throughout the entire model. We show that resulting dual embedding of sequence and structure in SSEmb is information-rich, leads to accurate predictions of variant effects probed by MAVEs, and can be used for other downstream tasks besides variant effect prediction. We exemplify this by using the SSEmb embeddings to predict protein-protein binding sites with results comparable to specialized state-of-the-art methods.

**Figure 1.**
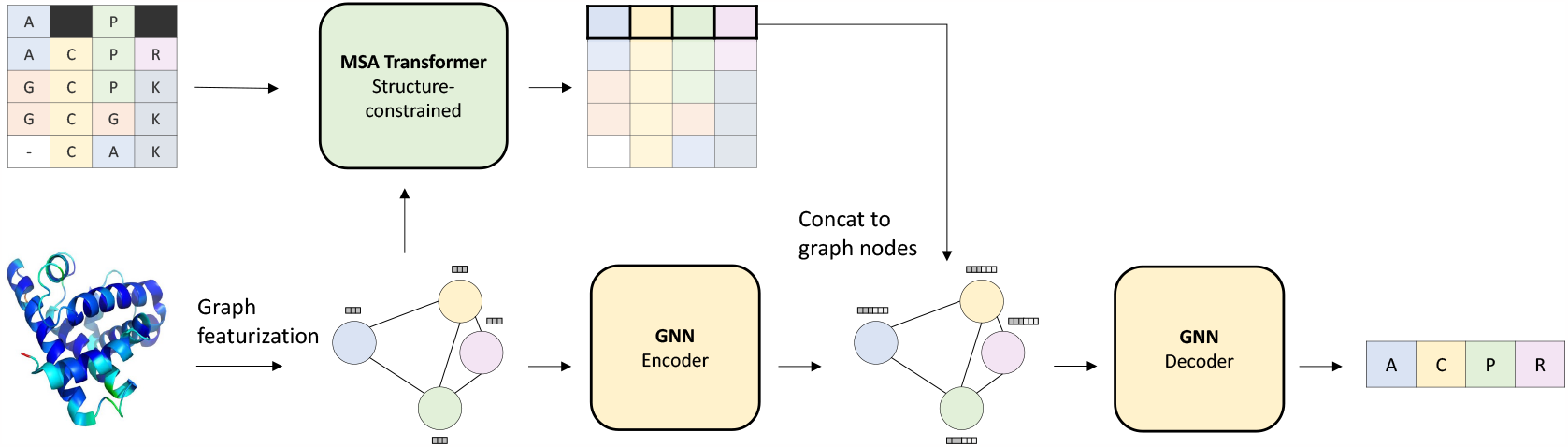
Overview of the SSEmb model and how it is trained. The model takes as input a subsampled MSA with a partially masked query sequence and a complete protein structure. The protein structure graph is used to mask (constrain) the row attention (i.e. attention across MSA columns) in the MSA Transformer. The MSA query sequence embeddings from the structure-constrained MSA Transformer are concatenated to the protein graph nodes. During training, SSEmb tries to predict the amino acid type at the masked positions. The model is optimized using the cross-entropy loss between the predicted and the true amino acid tokens at the masked positions. Variant effect prediction is made from these predictions as described in Methods.

## Results and Discussion

### Development of the SSEmb model

The SSEmb model was trained in a self-supervised manner using a combination of MSAs and protein structures (Fig. 1). The protein structures were taken from the previously compiled CATH 4.2 data set (***Ingraham et al., 2019***) that contains 18,204 training proteins with 40% non-redundancy partitioned by CATH class. For each of these protein structures we also generated a MSA (***Steinegger and Søding, 2017***). Before training, we removed proteins from the data set that were also present in the MAVE validation set or in the ProteinGym ***Notin et al. (2022a***) test set. During training, we mask a random subset of amino acid positions in the wild-type amino acid sequence, and the SSEmb model was trained to predict the masked amino acid type. We constrain the MSA Transformer with structure information by only allowing attention between positions in the MSA that are proximal in the three-dimensional protein structure. In order to combine information from the MSA and the protein structure at the model input level, features from the structure-constrained MSA Transformer were input to the nodes of the protein graph processed by the GNN module. Specifically, we extracted the last-layer embeddings from the structure-constrained MSA Transformer for each of the MSA query sequence positions and concatenated these embedding features to the nodes of the GNN. The structure-constrained MSA Transformer model was initialized using weights from the original pre-trained MSA Transformer, while the GNN was trained from scratch with an architecture similar to the Geometric Vector Perceptron (GVP) model (***Jing et al., 2021b***). Further details on SSEmb model architecture and training can be found in the Methods section.

### Validation using multiplexed assays of variant effects

During training, we validated the SSEmb model the results from ten MAVEs probing the effects of individual substitutions. Various model design choices along with relevant hyperparameters were selected based on SSEmb’s performance on these validation assays, which we selected based on two main criteria. First, the validation set should contain a mix of assays probing protein activity and abundance, with a majority of assays probing activity. Because protein activity and abundance are known to be correlated but distinct measures, this allowed us to obtain better insights into what the model was learning during training. Second, the validation set should contain a mix of assays considered either diffcult or relatively easier to predict with methods based either on protein structure or sequence alignments. Because these methods are known to capture some aspects of the data, this criterion enabled us to investigate whether the model was learning the same aspects as well as novel correlations not picked up by these methods. The assay data for the validation set was taken from ProteinGym (***Notin et al., 2022a***), with the exception of the LDLRAP1 assay where we used data from ***Jiang and Roth (2019); Livesey and Marsh (2023***) and the MAPK assay where we used data from ***Brenan et al. (2016); Livesey and Marsh (2020***).

To benchmark SSEmb performance during development, we compare our results to two non-machine-learning methods that use either only the MSA or only the protein structure as input. For the MSA-based model, we selected GEMME (***Laine et al., 2019***), which has been shown to produce state-of-the-art results for protein activity prediction outperforming most current machine learning methods using a relatively simple evolutionary model (***Notin et al., 2022a***). For the structure-based model, we selected stability predictions using Rosetta (***Park et al., 2016***), which are commonly used within protein engineering and have been shown to make useful predictions of protein stability and abundance (***Frenz et al., 2020; Høie et al., 2022; Gerasimavicius et al., 2023***).

After training, the SSEmb model achieves a higher Spearman correlation on the MAVE validation set than both GEMME and Rosetta (Table 1). In particular, the SSEmb model performs approximately the same as GEMME on the activity-based MAVEs that probe protein functions, but performs better overall on the abundance assays, suggesting that the added structural information in the SSEMb model helps it make better predictions for this problem. Overall, these results indicate that SSEmb is able to make accurate variant effect predictions that correlate well with measures of both activity and abundance.

**Table 1.** Overview of SSEmb results on the MAVE validation set after model training. We use the Spearman correlation coefficient to quantify the agreement between the data generated by the MAVEs and the predictions from SSEmb, GEMME and Rosetta. In this validation, only single-mutant variant effects were considered. The following protein structures were used in the SSEmb and Rosetta input: NUD15: 5BON_A, TPMT: 2H11_A, CP2C9: 1R90_A, P53: 4QO1_B, PABP: 1CVJ_G, SUMO1: 1WYW_B, RL401: 6NYO_E, PTEN: 1D5R_A, MAPK: 4QTA_A, LDLRAP1: 3SO6_A. The following UniProt IDs were used as input to construct multiple sequence alignments for GEMME: NUD15: Q9NV35, TPMT: P51580, CP2C9: P11712, P53: P04637, PABP: P11940, SUMO1: P63165, RL401: P0CH08, PTEN: P60484, MAPK1: P28482, LDLRAP1: Q5SW96.

| Protein | MAVE reference | MAVE type | Spearman $\vert \rho_S\vert $ | | |
| --- | --- | --- | --- | --- | --- |
|  |  |  | SSEmb | GEMME | Rosetta |
| NUD15 | Suiter et al 2020 | Abundance | **0.584** | 0.543 | 0.437 |
| TPMT | Matreyek et al 2018 | Abundance | 0.523 | **0.529** | 0.489 |
| CP2C9 | Amorosi et al 2021 | Abundance | **0.609** | 0.423 | 0.519 |
| P53 | Kotler et al 2018 | Competitive growth | 0.577 | **0.655** | 0.488 |
| PABP | Melamed et al 2013 | Competitive growth | **0.595** | 0.569 | 0.384 |
| SUMO1 | Weile et al 2017 | Competitive growth | **0.481** | 0.406 | 0.433 |
| RL401 | Roscoe & Bolon 2014 | E1 reactivity | **0.438** | 0.390 | 0.366 |
| PTEN | Mighell et al 2018 | Competitive growth | 0.422 | **0.532** | 0.423 |
| MAPK | Brenan et al 2016 | Competitive growth | 0.395 | **0.445** | 0.307 |
| LDLRAP1 | Jiang et al 2019 | Two-hybrid assay | **0.411** | 0.348 | 0.377 |
| Mean |  |  | **0.503** | 0.484 | 0.422 |

### Testing SSEMb on ProteinGym benchmark

ProteinGym is a large collection of data generated by MAVEs that has been used for benchmarking variant effect prediction models (***Notin et al., 2022a***). The ProteinGym substitution benchmark contains a total of 87 datasets on 72 different proteins. Of the 87 datasets in ProteinGym, 76 contain only single substitutions effects whereas the remaining 11 assays include variant effects from multiple substitutions. When testing the SSEmb model on the ProteinGym substitution set, we exclude a total of 9 MAVEs, which were part of our validation set. Furthermore, when multiple assays are present for a single UniProt ID, we report the mean correlation over the assays in order to remove potential biases from correlated measurements.

We compared the prediction accuracy of the trained SSEmb model the original MSA Transformer model on the ProteinGym set, and find improved accuracy (_*s*_ of 0.45 vs. 0.42) (Table 2 and Fig. S1). In line with the design of SSEmb, this difference is greatest for proteins with the shallowest MSAs (⟨ρ_*s*_ ⟩of 0.45 vs. 0.39) (Table 2 and Fig. 2). SSEmb also compares favourably to many other variant effect prediction methods benchmarked against ProteinGym (Table S1).

**Table 2.** Model performance on ProteinGym substitution benchmark grouped by UniProt ID and segmented by MSA depth. Low: *N*_eff_/*L <* 1, Medium: *N*_eff_/*L <* 100, High: *N*_eff_/*L >* 100 (***Notin et al., 2022a***).

| Model | Spearman $\rho_S$ by MSA depth | | | |
| --- | --- | --- | --- | --- |
|  | Low | Medium | High | All |
| SSEmb | **0.449** | **0.439** | **0.501** | **0.453** |
| MSA Transformer | 0.385 | 0.426 | 0.470 | 0.424 |

**Figure 2.**
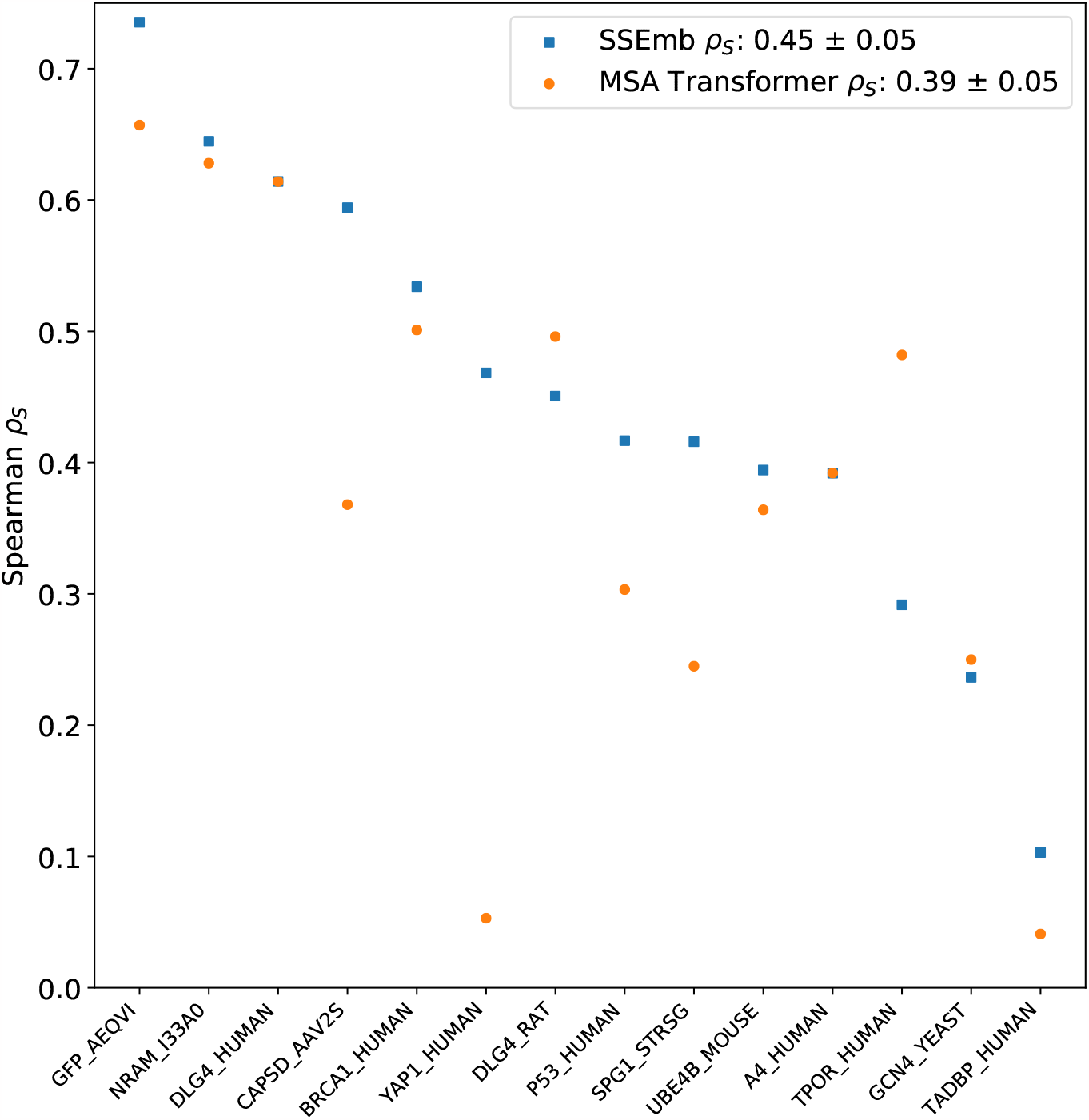
Overview of SSEmb results on the ProteinGym low-MSA (*N*_eff_/*L <* 1) substitution benchmark subset grouped by UniProt ID. Spearman correlations are plotted for both SSEmb (blue) and the MSA Transformer ensemble (orange). The mean and standard error of the mean of the set of all ProteinGym Spearman correlations are presented in the legend. Assays from the SSEmb validation set has been excluded from the original data set.

### Prediction of protein stability

Because of the tight interplay between protein stability, sequence conservation and cellular protein abundance (***Drummond et al., 2005; Yen et al., 2008; Serohijos et al., 2012; Matreyek et al., 2018; Suiter et al., 2020; Cagiada et al., 2021; Bédard et al., 2022***), we hypothesized that SSEmb would also be useful as a predictor of protein stability. We therefore tested the zero-shot performance of SSEmb on the recently described mega-scale measurements of protein stability (***Tsuboyama et al., 2023***). We find that SSEmb achieves an absolute Spearman correlation coefficient of 0.61 (Fig. S2), comparable to dedicated methods for protein stability predictions (***Blaabjerg et al., 2023***). This result demonstrates that SSEmb can be used as a zero-shot predictor of protein stability, and additional accuracy could possibly be achieved by further supervised learning (***Dieckhaus et al., 2023***).

### Prediction of protein-protein binding sites using embeddings

Protein-protein interactions (PPIs) are essential for cellular signaling and function. As such, mutations in PPI sites can often lead to disease (***Sahni et al., 2015; Mosca et al., 2015; Cheng et al., 2021***) and targeting protein-protein binding sites via pharmacological drugs is an ongoing area of research (***Winter et al., 2012; Scott et al., 2016***). Often, the protein-protein binding site consists of a small set of evolutionarily conversed surface residues that are essential for binding affinity (***Teppa et al., 2017***). Because these binding site residues are characterized by a combination of structural features (surface-proximity) and sequence features (evolutionary conservation), we hypothesized that the SSEmb model should contain some information relevant to the identification of these binding sites in its embeddings. Indeed, we have recently shown that a broader class of functional sites in proteins could be identified by a combined analysis of protein stability and conservation (***Cagiada et al., 2023***).

In order to test this hypothesis, we train a small supervised downstream model to predict protein binding sites from the SSEmb embeddings using PPI training and test data (***Tubiana et al., 2022***). Specifically, the downstream model takes as input the embeddings from the last layer in SSEmb and was trained to classify each residue as either belonging to a binding site or not using an attention mechanism (see Methods for further details). We compare our results to the state-of-the-art ScanNet model, which has been specifically developed for this classification task, as well as a xgboost baseline model with handcrafted structure- and sequence-based features (***Tubiana et al., 2022***). We evaluate the models using the area under the precision-recall curves (PR-AUC) and find that SSEmb downstream model performs in between the problem-specific ScanNet and baseline models across five different test sets with varying degrees of similarity to the training data (Table 3). These results demonstrate that the SSEmb embeddings contain a rich mix of structure- and sequence-based information, which may serve as useful features for binding site prediction and other downstream tasks.

**Table 3.** Using the SSEmb embeddings to study protein-protein interactions. The results show the PR-AUC for our supervised downstream model compared to ScanNet and a baseline model across five different test sets. All training and test sets as well as performance metrics for ScanNet and the handcrafted-features baseline model are from ***Tubiana et al. (2022***).

| Model | PR-AUC |  |  |  |  |
| --- | --- | --- | --- | --- | --- |
|  | Test set<br>(70%) | Test set<br>(homology) | Test set<br>(topology) | Test set<br>(none) | Test set<br>(all) |
| SSEmb downstream | 0.684 | 0.651 | 0.672 | 0.571 | 0.642 |
| Handcrafted features baseline | 0.596 | 0.567 | 0.568 | 0.432 | 0.537 |
| ScanNet | 0.732 | 0.712 | 0.735 | 0.605 | 0.694 |

## Conclusions

We have here presented a method for integrating information about protein sequence, conservation and structure in a single computational model. SSEmb uses a graph featurization of the protein structure both to constrain and integrate information from the corresponding MSA. Our results show that adding structural information to a pre-trained MSA-based model increases the ability of the model to predict variant effects in cases where the MSA is either lacking or shallow. We find that the embeddings learned by SSEmb during training contain information useful for downstream models. As an example, we show how a relatively simple downstream model trained with SSEmb embeddings as input is able to predict protein-protein binding sites at a level close to the state-of-the-art. We hope that SSEMb will serve as a useful tool for studying how the integration of sequence- and structure-based protein information can improve computational predictions of variant effects, and could for example be used to disentangle mechanistic aspects of variant effects (***Cagiada et al., 2023***).

## Methods

### Multiple sequence alignments

MSAs were generated using MMSeqs2 (***Steinegger and Søding, 2017***) in combination with filtering via sequence identity buckets as implemented in ColabFold (***Mirdita et al., 2022***), which aims to maximizes the diversity of sequences in the final alignment. This protocol to generate MSAs has been shown to work well for variation effect prediction using the GEMME model (***Abakarova et al., 2022***). We use the original ColabFold Search protocol with the parameters `–-diff=512`, `–-filter-min-enable=64`, `–-max-seq-id=0.90`. Furthermore, we add the parameter `–-cov=0.75` to each sequence indentity bucket in order to ensure that we only retrieve high-coverage sequences for the generated MSAs.

### Subsampling of multiple sequence alignments

We randomly subsample the full MSA before using it as input to SSEmb in order to make the model train with GPU memory constraints. During training, the number of subsampled MSA sequences is set to 16. We explored how variant effect prediction performance on the validation set scales with MSA subsampling depth and ensembling. We find SSEmb is more robust to MSA sequence depth compared to the original MSA Transformer (***Meier et al., 2021***). Furthermore, we find that ensembles of shallow MSAs out-perform single-use or ensembles of deeper MSAs (Fig. S3). In our case, ensembles are created by subsamling multiple times and taking the mean over the final variant effect predictions. Based on these results, we used 16 subsamled MSA sequences in an ensemble of 5 during model inference.

### Structure-constrained MSA Transformer

The structure-constrained MSA Transformer model used in SSEmb is based on the original architecture (***Rao et al., 2021***). At initialization, we use the pre-trained model weights (https://github.com/facebookresearch/esm/tree/main/esm). We modify the original MSA Transformer by applying a binary contact mask to the attention maps going across MSA columns (i.e. row attention) before normalization. The contact mask corresponds to the 20 nearest neighbour graph structure used in the SSEmb GNN module, ensuring that row attention values are only propagated for positions that are spatially close in the three-dimensional protein structure. During training, we only fine-tune the row attention layers of the structure-constrained MSA Transformer in order to conserve the phylogenetic information encoded in the column attention layers (***Lupo et al., 2022***).

### GNN module

The SSEmb GNN module follows the architecture of the GVP model (***Jing et al., 2021b***), with a few important adjustments: (i) Graph edges were defined for the 20 closest nodes neighbors instead of 30 in the original implementation, (ii) Node embedding dimensions were increased to 256 and 64 dimensions for the scalar and vector channels respectively. Edge embedding dimensions were kept at 32 and 1 as in the original implementation, (iii)The number of encoder and decoder layers were increased to four, (iv) we used vector gating (***Jing et al., 2021a***), (v) the MSA query sequence embeddings from the structure-constrained MSA Transformer were concatenated to the node embeddings of the GNN decoder and passed through a dense layer to reduce dimensionality, and (vi) the models prediction task was changed from auto-regressive sequence prediction to a masked token prediction task. Further details are described below in the ‘Model training’ section.

### Model training

During SSEmb training, we randomly mask amino acids in the wild-type sequence following a modified BERT masking scheme (***Devlin et al., 2019***). Before each forward pass, 15% of all wild-type sequence residues are selected for optimization. Out of these 15% residues, 60% are masked, 20% are masked together with all residues in the corresponding MSA column are masked, 10% are replaced by a random residue type, and 10% are left unchanged. After masking, the SSEmb model is tasked to predict the amino acid types of the masked residues, given the protein structure and a subsampled MSA input. The masked prediction task is optimized using the cross-entropy loss between the selected 15% wild-type amino acid types and the corresponding predicted amino acid types. We trained SSEmb using the gradual unfreezing method in two steps (***Howard and Ruder, 2018***). First, we trained the GNN module until the model was close to convergence, while keeping the parameters in the structure-constrained MSA Transformer frozen. Second, we unfreeze the row attention parameters in the structure-constrained MSA Transformer and fine-tune both the GNN module and the structure-constrained MSA Transformer until convergence. Training was performed using the Adam optimizer (***Kingma and Ba, 2014***) with a learning rate of 10^−3^ for the GNN module and 10^−6^ for the structure-constrained MSA Transformer respectively. Batch sizes were fixed at 128 and 2048 proteins for the two training stages respectively.

### Variant effect prediction

At inference, we randomly subsample 16 sequences from the full MSA five times with replacement in order to generate an ensemble of model predictions. The final SSEmb score is computed as the mean of the scores from this ensemble. Protein variants scores were computed according to the masked marginal method (***Meier et al., 2021***) from:

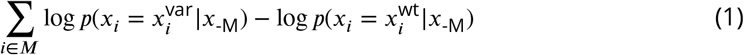

where *x*^var^ and *x*^wt^ represent the variant (mutant) and wild-type sequences, and *x*_-M_ represent a sequence where the set of substituted (mutated) positions *M* have been filled with mask tokens. The above model represents an additive variant effect model.

### Predicted protein structures for ProteinGym test

We used AlphaFold (***Jumper et al., 2021***) to predict structures for the proteins in the ProteinGym substitution benchmark. In practice, we used the ColabFold implementation (***Mirdita et al., 2022***) with default settings. For each sequence, we selected the predicted protein structure with the highest rank as input to SSEmb.

All protein structures used as input to the SSEmb model during testing were preprocessed using the OpenMM PDBFixer package (***Eastman et al., 2013***). The protein structures in the training set were used without modifications.

### Rosetta protocol

Rosetta ΔΔ*G* values were computed using the Cartesian ΔΔ*G* protocol (***Park et al., 2016***) and the Rosetta version with GitHub SHA1 224ebc0d2d0677ccdd9af42a54461d09367b65b3. Thermodynamic stability changes in Rosetta Energy Units were converted to a scale corresponding to kcal/mol by dividing with 2.9 (***Park et al., 2016***).

### GEMME protocol

GEMME scores were computed using default settings (***Laine et al., 2019***). MSAs for the GEMME input were generated as previously described (***Høie et al., 2022***) using Hhblits version 2.0.15 (***Remmert et al., 2011***) to search the UniRef30 database with settings: `-e 1e-10 -i 1 -p 40 -b 1 -B 20000`. We applied two additional filters to the HHblits output MSA before using them as input to GEMME. The first filter removes all the positions (columns) that are not present in the query sequence and the second filter removes all the sequences (rows) where the number of total gaps exceeds 50%. We note that other ways of constructing MSAs may improve the accuracy of GEMME (***Abakarova et al., 2022***); in the comparison to the ProteinGym benchmark (Table S1) we therefore used data directly from ProteinGym.

### Filtering of mega-scale protein stability data set

We tested the accuracy of of SSEmb in zero-shot predictions of changes in protein stability using data set 3 from ***Tsuboyama et al. (2023***), which consists of experimentally well-defined ΔΔ*G* measurements for a total of 607,839 protein sequences. The data set was used with minor filtering. First, our model is focused on predicting the effects of non-synonymous mutations and we therefore removed all synonymous, insertion and deletion mutations. Second, we discarded a total of 75 protein domains for which no corresponding AlphaFold model was included in the original data set.

### Downstream model for protein-protein binding site prediction

We used the PPBS data set (***Tubiana et al., 2022***) for protein-protein binding site predictions. The original data set contains a total of 20,025 protein chains with binary residue-level binding site labels. We filtered the data set to exclude proteins chains that were marked as obsolete in the RCSB Protein Data Bank, where the protein chain had missing binding site labels or where the amino acid sequence in the structure did not match the sequence in the label data. Lastly, we removed protein chains that were longer than the 1024 amino acid sequence limit imposed by the MSA Transformer. The total number of protein chains in the modified PPBS data set is 19,264.

The SSEmb downstream model consisted of a small Transformer encoder model, which follows the original encoder implementation (***Vaswani et al., 2017***). The down-stream model takes as input the last-layer 256-dimensional residue-level embedding vector from the SSEmb model as the only input. In order to compute all embeddings in a reasonable time frame, we did not mask any positions in the amino acid sequence inspired by the wild-type marginal method (***Meier et al., 2021***):

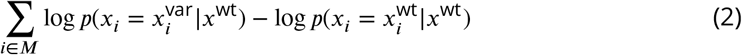

This method is much faster than the masked marginal method as it only needs a single forward pass to compute embeddings for all positions in the sequence and it has been shown to approximate the masked marginal method well for variant effect prediction (***Meier et al., 2021***). Positional encodings are added using sine and cosine functions and attention is applied across the entire amino acid sequence. The number of hidden dimensions is kept fixed at 256 and the sequence is processed using three attention layers with three attention heads each. The downstream model is optimized in a supervised manner on the training data (***Tubiana et al., 2022***) using a binary cross entropy loss and the Adam optimizer (***Kingma and Ba, 2014***) with a learning rate of 10^−4^ and a batch size of ten proteins. The accuracy of the final model was evaluated using the area under the precision recall curves (Table 3 and Fig. S4)

## Data and code availability

Scripts and data to repeat our analyses are available via https://github.com/KULL-Centre/_2023_Blaabjerg_SSEmb.

## Acknowledgement

Our research is supported by the PRISM (Protein Interactions and Stability in Medicine and Genomics) centre funded by the Novo Nordisk Foundation (NNF18OC0033950 to A.S. and K.L.-L.), and by grants from the Carlsberg Foundation (CF21-0392 to K.L.-L.), Novo Nordisk Foundation (NNF20OC0062606 and NNF18OC0052719 to W.B.) and the Lund-beck Foundation (R272-2017-4528 to A.S.).

