## Supporting Figures and Table for "A joint embedding of protein sequence and structure enables robust variant effect predictions"

### Supporting material for: A joint embedding of protein sequence and structure enables robust variant effect predictions

| Model | Spearman $\rho_S$ by MSA depth | | | |
| --- | --- | --- | --- | --- |
| | Low | Medium | High | All |
| TranceptEVE L | **0.451** | **0.462** | 0.502 | **0.468** |
| GEMME | 0.429 | 0.448 | 0.495 | 0.453 |
| SSEmb | 0.449 | 0.439 | 0.501 | 0.453 |
| Tranception L | 0.438 | 0.438 | 0.467 | 0.444 |
| EVE (ensemble) | 0.412 | 0.438 | 0.493 | 0.443 |
| VESPA | 0.411 | 0.422 | **0.514** | 0.438 |
| EVE (single) | 0.405 | 0.431 | 0.488 | 0.437 |
| MSA Transformer (ensemble) | 0.385 | 0.426 | 0.470 | 0.426 |

**Table S1.** Model performance on ProteinGym substitution benchmark compared to other variant effect prediction models grouped by UniProt ID and segmented by MSA depth. Low: $N_{\text{eff}}/L < 1$, Medium: $N_{\text{eff}}/L < 100$, High: $N_{\text{eff}}/L > 100$ (***Notin et al., 2022***).

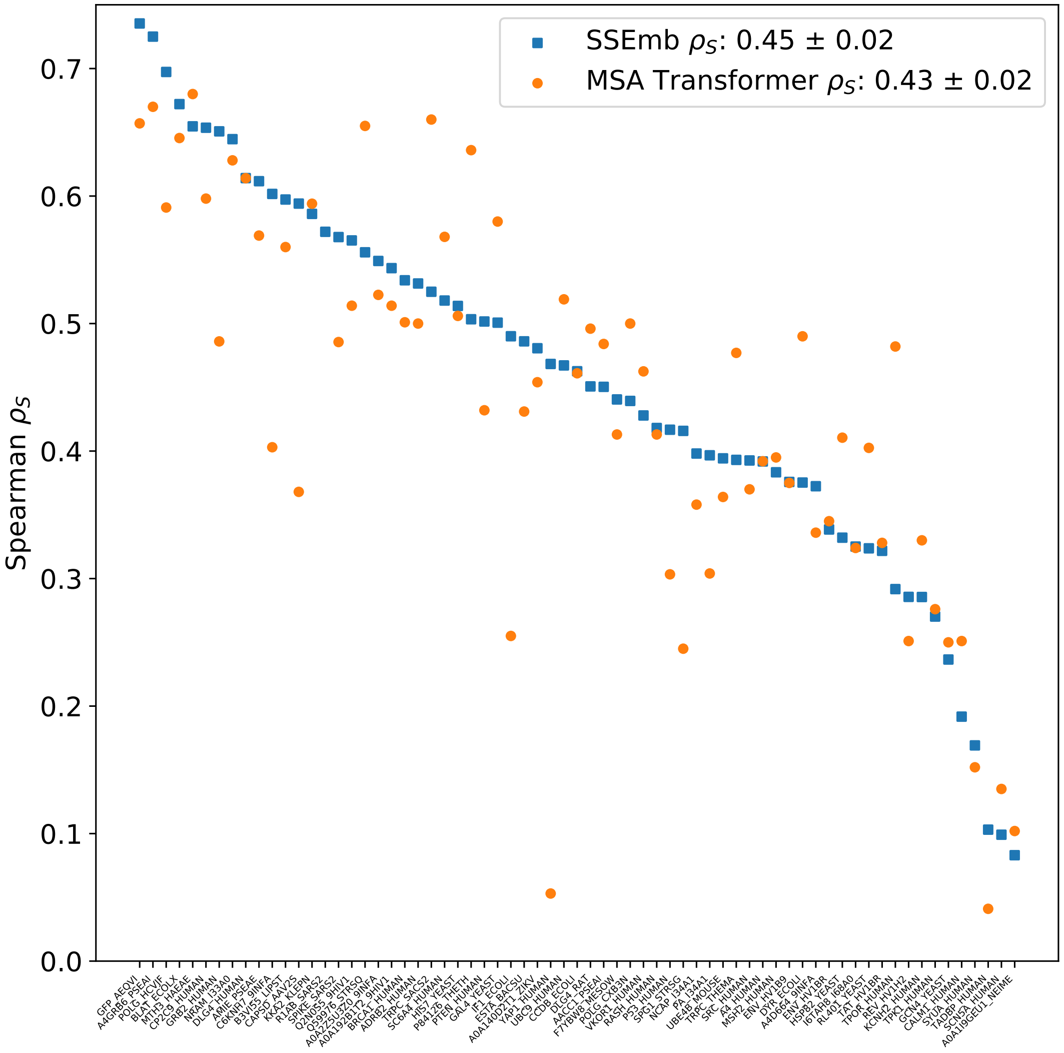


**Figure S1.** Overview of SSEmb results on the full ProteinGym substitution benchmark grouped by UniProt ID. Spearman correlations are plotted for both SSEmb (blue) and the MSA Transformer ensemble (orange). The mean and standard error of the mean of the set of all ProteinGym Spearman correlations are presented in the legend. Assays from the SSEmb validation set have been excluded from the original data set.

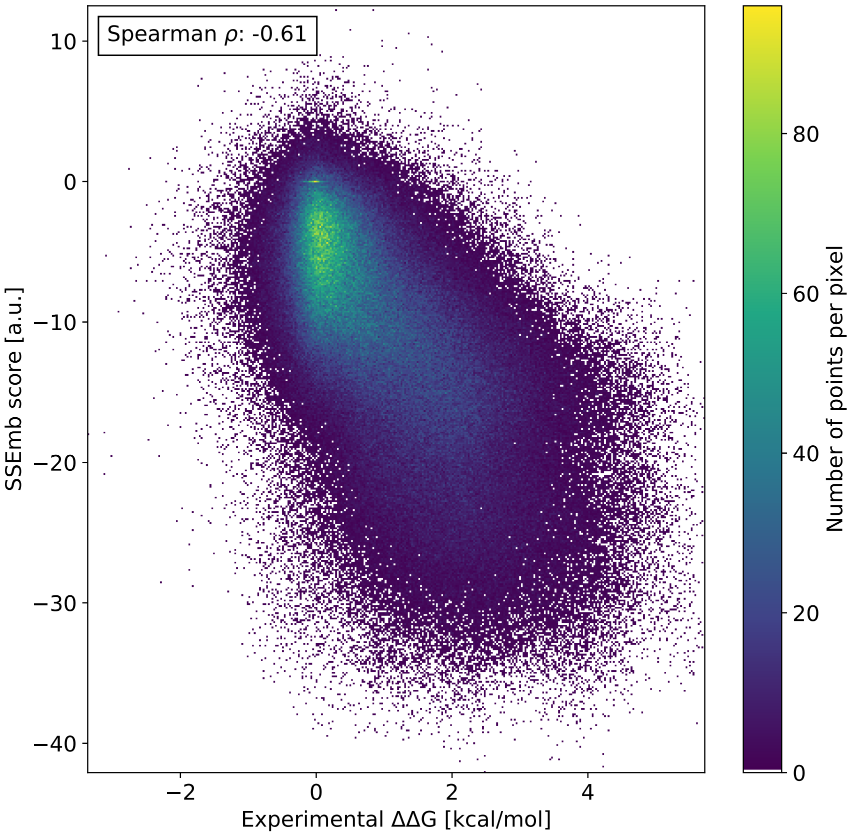


**Figure S2.** SSEmb scores versus experimental $\Delta\Delta G$s from the recently published mega-scale experiments (***Tsuboyama et al., 2023***). The experimental data set was filtered to include only substitution variants and variants with a corresponding structural model from AlphaFold.

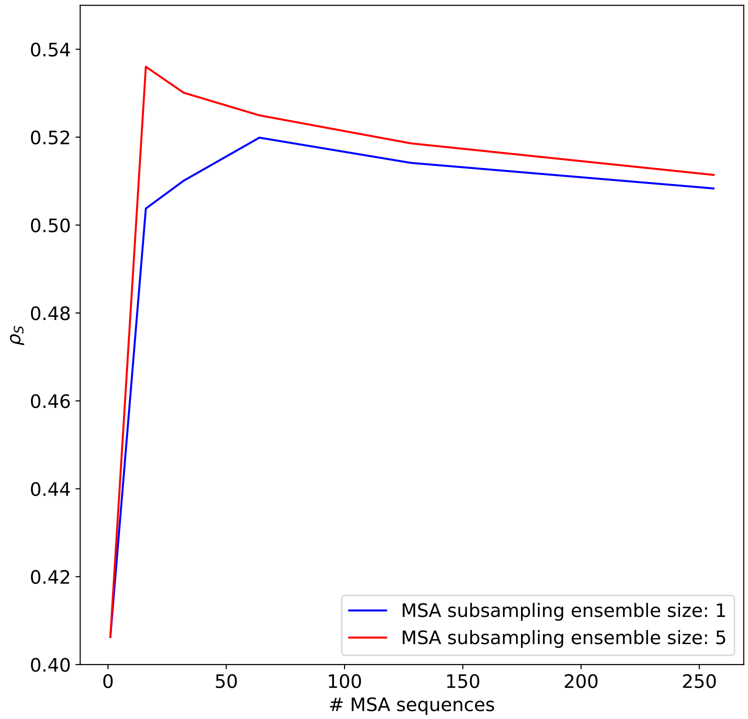


**Figure S3.** Mean Spearman correlation between SSEmb predictions and MAVE scores for ten validation proteins as a function of the number of subsampled MSA sequences used as SSEmb input. SSEmb is robust to the number of input MSA sequences reaching optimal performance at approximately 16 MSA sequences. SSEmb benefits from ensembles of randomly subsampled MSA sequences with the effect gradually decreasing as the number of subsampled sequences approaches the total number of sequences in the full MSA.

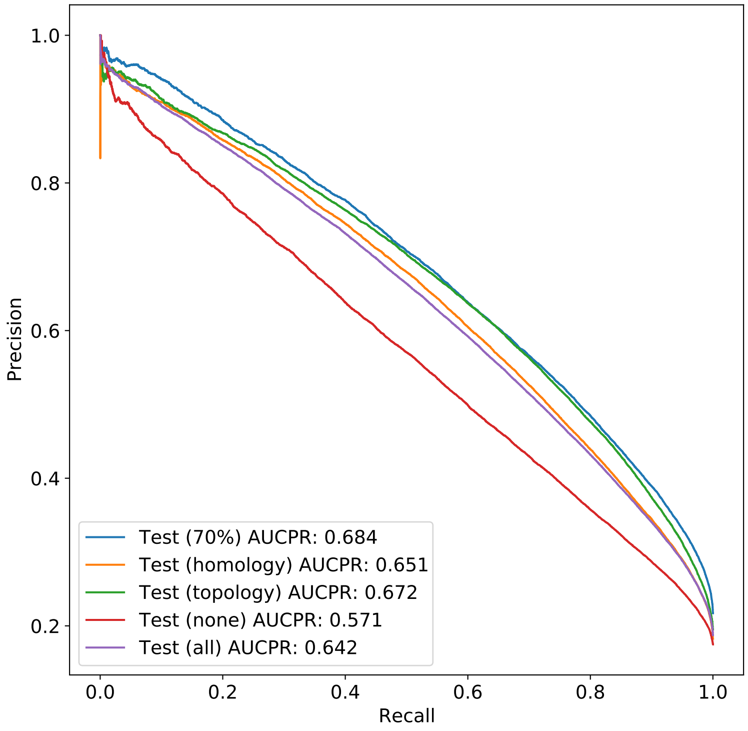


**Figure S4.** Precision-recall curves for the downstream model trained to predict protein-protein binding sites using the PPBS data set (***Tubiana et al., 2022***). Curves for different test sets with varying degrees of similarity to the training data is shown.
